# Reciprocal E-cadherin signaling aligns apical surfaces between neighboring epithelial tissues to complete the *C. elegans* digestive tract

**DOI:** 10.1101/2025.10.06.680605

**Authors:** Lauren E. Cote, Maria D. Sallee, Rachel K. Ng, Melissa A. Pickett, Jérémy Magescas, Jessica L. Feldman

**Author notes:** College of Idaho, Caldwell, ID, United States.

## Abstract

Epithelial cells form barriers with specialized apical membranes facing the external environment or internal lumens, yet how interconnected epithelial tubular systems form from cystic epithelial primordia is poorly studied. At the boundaries of cystic epithelial primordia, apical surfaces must correctly align to yield functional connections within and between organs. Here, we use the digestive tract of the developing *C. elegans* embryo to define a crucial two-cell tissue, the rectal valve (vir) cells, that mediate the connection and apical alignment between the posterior intestine and the rectum. Vir cells migrate while attached to the rectal primordium to create an essential connection with the intestinal primordium and complete the digestive tract. Vir contact with the intestine, through E-cadherin signaling, is required for the intestinal apical surface to undergo a transition from a cyst with an inaccessible apical surface to an open tube competent to connect with neighboring tissues. Ablation of vir cell progenitors results in a lethal disconnection between the intestinal and rectal apical surfaces due to failure of this first step in apical accessibility. To resolve the final gap in apical continuity between the intestine and rectum, vir cells develop bipolar apical surfaces, each facing the neighboring tissue. The intestine-facing apical puncta in vir require intestinal E-cadherin signaling to form and accumulate as well as to correctly partition apical proteins within vir cells. Together these results establish E-cadherin-based signaling as the crucial factor aligning apical surfaces across primordia as epithelial cells from different germ layers form a vital connection.

## Results

The most common body plan in the animal kingdom is an ‘epithelial toroid’ which involves epithelial cells from endoderm and ectoderm connecting^1,2^. These epithelial cells have specialized apical membranes that face internal lumens or the external environment^3–8^ and readily build individual closed-off cysts in developing primordia or in in vitro culture models. Yet, how enclosed cysts connect to form functional interconnected tubular systems such as sensory, reproductive, excretory, circulatory, and digestive systems is poorly understood^9–14^. At the cyst-cyst interface between epithelia primordia, cells that may have different developmental origins, polarization mechanisms, and functions must nonetheless align to create a single continuous apical surface. Ectopic or inappropriate connections can lead to fistulas while absence of correct apical alignment between fusing epithelial structures can lead to congenital atresias and other abnormalities^15–19^. Here, we use the development of the C. *elegans* digestive tract to investigate the alignment of apical surfaces across tissues, focusing on the final connection between the intestine and rectum that completes the epithelial toroid (Figure 1A, B).

**Figure 1:**
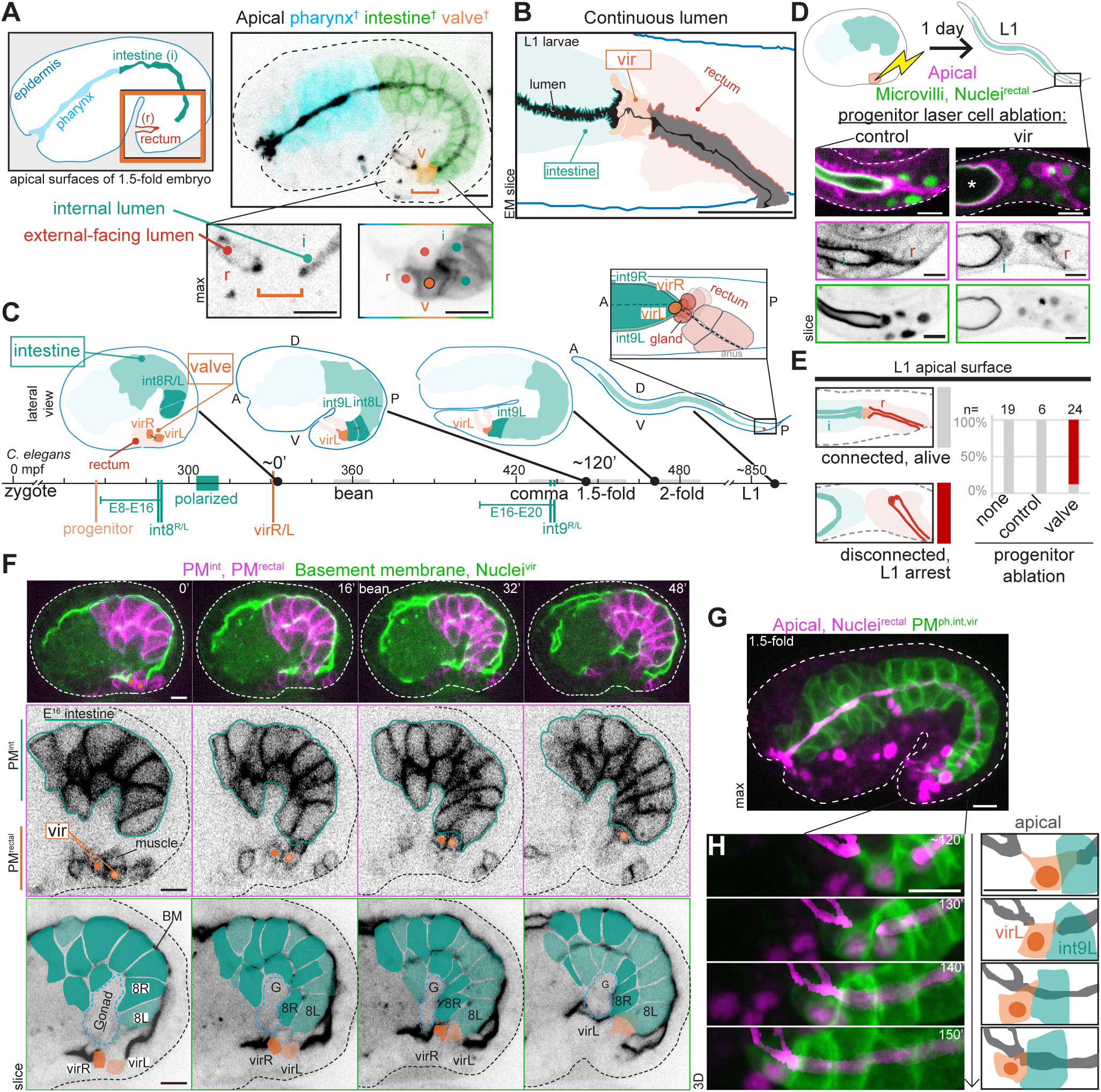
Rectal valve cells (vir) are a two-cell tissue that mediates the intestine-rectal connection. A) Live image of 1.5-fold stage embryo expressing endogenously-tagged PKC-3 (black, “Apical” throughout) and digestive tract <u>p</u>lasma <u>m</u>embrane (*pha-4p*::PM recolored by tissue^†^). **<u>v</u>**alve-intestinal-rectal (vir, orange throughout) cells localize within final apical discontinuity between the **<u>r</u>**ectum and **i**ntestine (teal throughout). Orange box, inset location in all figures unless noted, depicting orientation (rectum on left). B) Tracing of continuous larval lumen (black) from serial section EM^27^. C) Morphogenetic timeline of posterior intestine and vir. mpf: minutes post fertilization with timing relative to vir birth used in all timelapses. Vertical lines: cell divisions. E8-E16-E20: intestinal ‘E’ cell number. D, E) Larval intestine-rectal connection (PKC-3, magenta; EPS-8A, *ceh-27p*::<u>H</u>istoneH1, green) after embryonic progenitor laser cell ablation. *, intestinal cyst. F) Timelapse of intestinal (*end-1p*::PM, magenta) and virR/L (*nhr-67p*::PM, magenta; FOS-1A, green nuclei) membranes and basement membrane (Laminin-2, green). G, H) Timelapse imaging of merging apical surfaces (PKC-3, magenta, highlighted in H (Methods)) within the digestive tract (*pha-4p*::PM, green) during embryonic elongation with vir marker (*nhr-67p*::H1, magenta). Cartoons in H aligned to rectum. All figures unless noted: live confocal image of *C. elegans* embryo in lateral view expressing endogenous fluorescently-tagged protein(s) (‘max’: maximum intensity projection through tissue; ‘slice’: single z-slice; ‘3D’: 3-dimensional napari rendering). Single channel images boxed in corresponding color. Scale bars, 5 µm.

### Rectal valve cells (vir) mediate the intestinal-rectal connection

To determine how the continuous apical surface of the epithelial toroid develops, we explored the endogenous localization of the fluorescently-tagged apical protein aPKC/PKC-3. We found a continuous line of apical proteins from the skin through the pharynx to the intestine, but we observed a break in the continuity of the apical surface between the intestinal primordium and the invaginating rectal lumen at the posterior of the embryo indicating that this surface was the final connection completing the epithelial toroid (Figure 1A, n = 303/305 1.5-fold embryos). At this 1.5-fold stage of the embryo, both the intestine and rectum were polarized with apical and junctional proteins concentrated towards the future site of the lumen^4,20–26^, yet these apical surfaces were separated by a pair of cells sitting at their interface that corresponded to the rectal valve cells (‘vir’, <u>v</u>alve-intestinal-rectal: virR and virL, orange) (Figure 1C, Figure S1A-E). In serial section electron microscopy (EM) of larvae^27^ that shows the final form of the digestive tract, the ∼2um long interface between vir cells contains an extremely thin (50-65nm wide) and electron-dense sieve between the intestinal and rectal lumens (Figure 1B, Figure S1F) to ensure that the intestine only opens to the rectum during waste expulsion^28^. Ablation of the vir progenitor before vir-intestinal contact resulted in a disconnection between the intestinal and the rectal lumens^29^ (Figure 1D,E), although vir cells were dispensable for rectal invagination (Figure S1G). The resulting ‘blind gut’ caused arrest at the L1 larval stage, consistent with genetic evidence that rectal deformities are associated with lethality^30–33^, presumably due to the inability of the worms to expel waste. Therefore, vir cells provide a specific and essential connection that completes the digestive tract.

To investigate the morphogenesis of the rectal valve, we first observed that the two vir cells arise from a single progenitor that is lineally and spatially distinct from the intestinal progenitor (Figure 1C, Figure S2A-G, Video S1). The progenitors of vir and the 9 other rectal cells are born on the ventral surface and all arise from the AB-lineage^29^. At this stage of development, the E blastomere-derived intestine is composed of 16 cells (‘E16’) that are named according to which of the nine intestinal rings (int1-int9) they give rise to^34^. Live imaging revealed that the vir progenitor gastrulated with its non-rectal neighbors to internalize^35^ (Figure S2F), divided before the remaining rectal cells invaginated, and was initially separated from the intestine by muscle and germ cells^36^ (Figure 1F). The vir cells contacted the intestinal primordium (Figure 1C,F, Figure S2C-E, Video S1) after the intestine had polarized^37^ (Figure S2C) and well before the final intestinal divisions (‘E16-E20’, Figure S2E). Whole-embryo cell membrane segmentation^36^ showed that virL and virR stably contacted the posterior-most intestinal cells (int8L and in8R) ∼15 min after vir cell birth (Figure S2B, Video S1) during the intercalation of int5 cells within the intestinal primordium^34^ (Figure S2D), although transient interactions between the progenitor and/or the posterior virL sometimes occurred. The more anterior virR cell then rotated to form a bilaterally symmetric interaction with the posterior end of the intestine (Figure S2G). At the 1.5-fold stage, about 2 hours after the initial contact between the vir cells and the posterior intestine, the break in the apical surface (Figure 1A) started to close, with the intestinal apical surface appearing to extend towards and join with the apical surface of the rectum during elongation (Figure 1G,H, Figure S2I-K, Video 2).

Specialized cells often breach a basement membrane to facilitate the joining of tissues, as in the *C. elegans* vulva, mammalian mouth, and kidney^38–41^. However, we found no basement membrane between vir and the posterior end of the intestine and instead the basement membrane surrounded the digestive tract after the vir cells made stable contact with the intestine (Figure 1F, Figure S2H). Therefore, the vir cells form a specialized tissue that bridges between the internal intestinal lumen and the external-facing rectal lumen through cell-cell contact.

### The rectal valve cells are required for a cyst-to-tube transition in the intestine

Given that larvae after vir precursor ablation showed an abrupt end to the intestinal lumen (Figure 1D, Figure S1G), we next explored how the apical surface of the intestine normally accesses the rectum during morphogenesis. Imaging from the ventral surface during the early E16 stage showed that the intestinal apical surface was present at the time when the vir cells made contact but did not reach the end of int8L/R (Figure S2C), forming a closed off cystic apical surface that was inaccessible to other cells. Shortly after the valve cells contacted the intestine, the intestinal apical surface appeared to extend and erode the cell-cell interface between the posterior int8L and int8R cells, converting the cyst into an accessible tube by redistributing apical signal (Figure 2A,A’,B, Figure S3A-C, Video 3). To test whether interaction with vir cells was required for the intestine to undergo this cyst-to-tube transition, we examined embryos at late bean stage, the time when the cyst-to-tube transition is complete, after laser cell ablation of the vir progenitor. Control ablations of cells neighboring the vir progenitor at the ventral midline resulted in a normal cyst-to-tube transition, indicating that thermal damage did not delay or block this apical extension within the intestinal cells (Figure 2C,D, Figure S3D-E). In contrast, valve progenitor ablation blocked the cyst-to-tube transition, indicating that the ‘blind gut’ visible in larvae (Figure 1D,E, Figure S1G) was a result of a failure of the first step in joining the intestine lumen to the rectum: creating an accessible apical surface at the end of the intestine.

**Figure 2:**
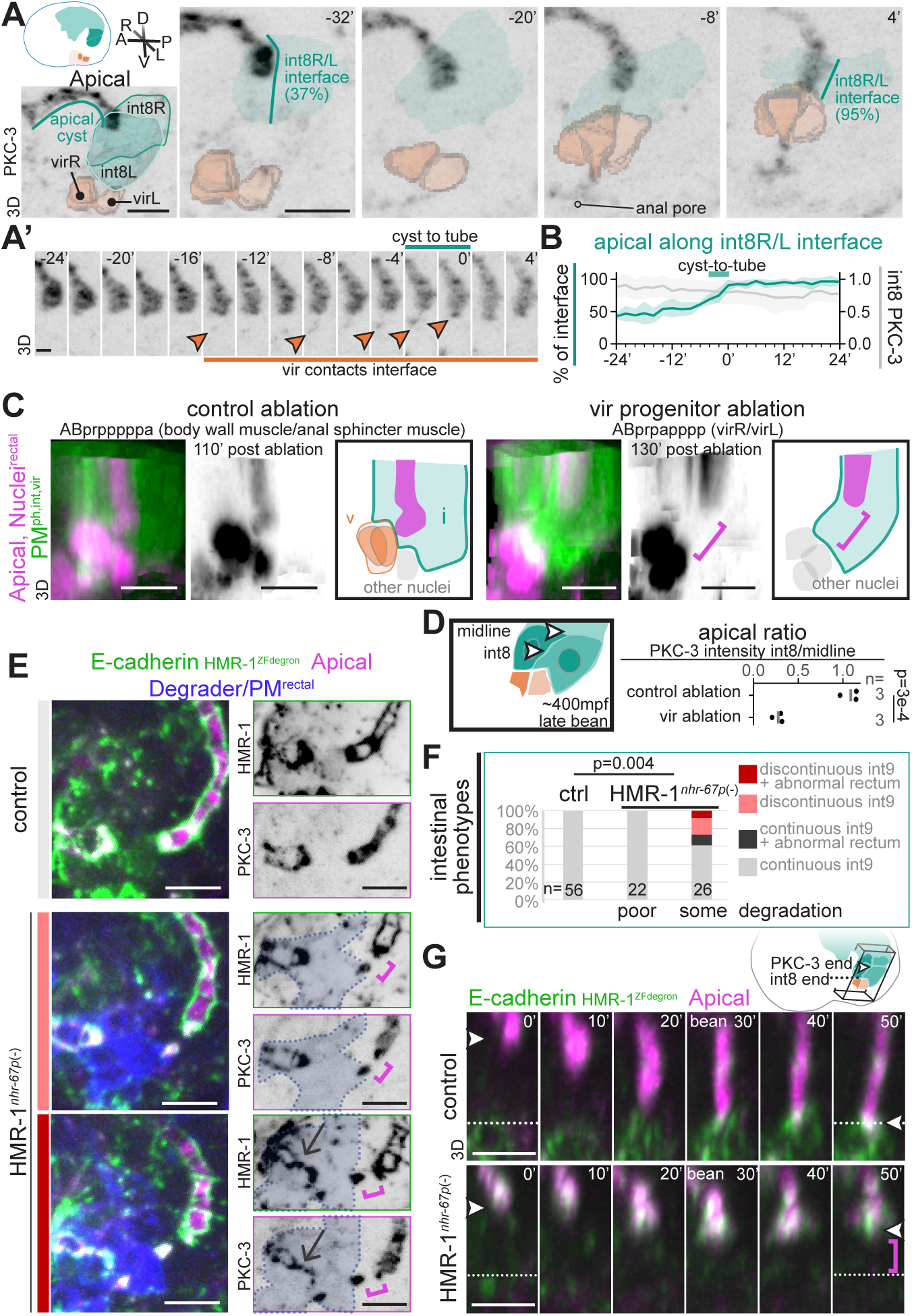
Rectal valve cells induce a cyst-to-tube transition in the intestine mediated by E-cadherin. A) Ventrolateral 3D renderings of apical PKC-3 time course along the int8L/R interface (light green) with 3D cell reconstructions from *pha-4p*::PM. A’) Appearance of small puncta (orange arrowhead) at the valve/intestinal contact. scale bar, 2 µm. B) Percentage coverage of the int8R/L midline interface by PKC-3 plotted over time (teal, mean and standard deviation of n=4 embryos where t=0 is completion of the cyst to tube transition) and total, bleach corrected PKC-3 intensity across int8 normalized to intensity at the initial timepoint imaged (grey). C) Cyst-to-tube transition (PKC-3, magenta, gamma=0.5 for simultaneous apical and nuclei visualization) in intestine (*pha-4p*::PM, green) after laser ablation of indicated progenitor (*nhr-67p*::H1, magenta, gamma=0.5). D) Normalized int8 PKC-3 intensity plotted as mean (p-value, t-test). E,F) Degradation of ZF degron-tagged E-cadherin (HMR-1, green) in *nhr-67*+ cells (blue (dotted outline)) shown in maximum intensity projection. Magenta bracket, apical gaps in PKC-3 (magenta). Arrow, abnormal rectum. F) Intestinal apical gaps binned by degradation strength (Figure S3I). p value=0.004, Fisher’s Exact Test. G) Cyst-to-tube transition time courses. Apical surface did not extend in n=0/8 control and n=2/6 HMR-1*^nhr-^*^67p^(-) embryos. Controls in E-G lack degradation transgene. All images unless noted are lateral 3D renderings. Scale bars, 5 µm.

Next, we wanted to test the role of cell-cell contact between the vir cells and the intestine in guiding the localization of apical proteins during the cyst-to-tube transition. The homophilic transmembrane adhesion protein E-cadherin/HMR-1 underlies cell-cell contact in many types of epithelia^42,43^, In *C. elegans*, the E-cadherin homolog HMR-1 is expressed during pharyngeal valve intercalation^44^, required to align apical surfaces along the polarizing intestine^45^, and zygotic HMR-1 mutants occasionally show a disconnection between the pharynx and the epidermis^46^. We therefore tested whether E-cadherin in the vir cells is required during vir cell contact with the intestine to drive the intestinal cyst-to-tube transition. To degrade E-cadherin in vir cells prior to their contact with the intestine, we used a tissue specific degradation system in which the target is endogenously tagged with a ZF degron and the degrader protein ZIF-1 is expressed in a cell or tissue of interest^47,48^. We used a ZF-tagged allele of HMR-1^45,49^ and expressed ZIF-1 with the promoter for *nhr-67* (*nhr-67*p, HMR-1^nhr-67^(–)), a gene which is expressed in the vir precursor cells as well as other ABp-derived cells^50,51^ (Figure S3F,G). Although this approach yielded incomplete HMR-1 depletion, a small but consistent minority of 1.5-fold stage HMR-1^nhr-67^(–) embryos showed intestinal defects (Figure 2E,F, Figure S3H-I). While the intestinal primordium was generally polarized in HMR-1^nhr-67^(–) embryos, we observed breaks in the apical surface at the posterior end of the intestine, suggesting a potential failure in the cyst-to-tube transition. Live imaging of this process in HMR-1^nhr-67^(–) embryos revealed an inability of apical proteins to spread between the left-right int8 surfaces with one showing erroneous spreading of the apical surface along the orthogonal surfaces between int7 and int8 (Figure 2G). These observations are consistent with a role for E-cadherin within vir cells to signal the reorientation of the apical surface in intestinal cells that underlies the cyst-to-tube transition. Together, these data show that vir cell contact is required for the first step in joining the intestinal lumen to that of the rectum: extending and exposing the apical surface at the posterior end of the intestine.

### Rectal valve cells are bipolar

Once the apical surface of the intestine is exposed and therefore competent to join with the apical surface of the neighboring tissues, the next morphogenetic step is to align and connect the rectal and intestinal apical surfaces (Figure 1H). We therefore next explored how vir cells join the end of the intestinal apical surface to the tip of the rectal lumen. Live imaging showed large puncta of apical and junctional proteins at the intestine-valve and valve-rectum interfaces (Figure 1H, Figure S2K) that merged during embryonic elongation from the 1.5-to 2-fold stage, filling in the hole in apical continuity. Due to the resolution limits of our imaging, we were unable to determine whether these endogenous puncta were wholly, partially, or not at all localized within the vir cells as opposed to within the neighboring intestinal and/or rectal tissues (Figure 1A). We therefore investigated if vir cells themselves contribute to this punctate apical signal by creating PAR-6^vir^, an apical reporter inserted into the valve-specific endogenous *fos-1a* locus^50,52^ (Figure S4A-B, Methods). PAR-6^vir^ localized to an intestine-facing punctum and a rectal-facing punctum (black and white arrowheads, respectively, Figure 3A-E, Figure S4C-E) and overlapped with endogenous PKC-3 signal (Figure 3A’). Super-resolution imaging of embryos at this stage showed multiple substructures within the intestine-facing and rectum-facing puncta (Figure 3D) that likely correspond to bipolar and not antipolar apical surfaces from each vir cell. Additionally, EM of 3-fold embryos^53^ showed that vir cells have contralateral contacts with both intestinal cells and that both vir cells contact the rectal lumen (Figure S1F, Figure S4F,G). These results are consistent with both vir cells having bipolar localization of apical proteins prior to the completion of apical connectivity (Figure 3E).

**Figure 3:**
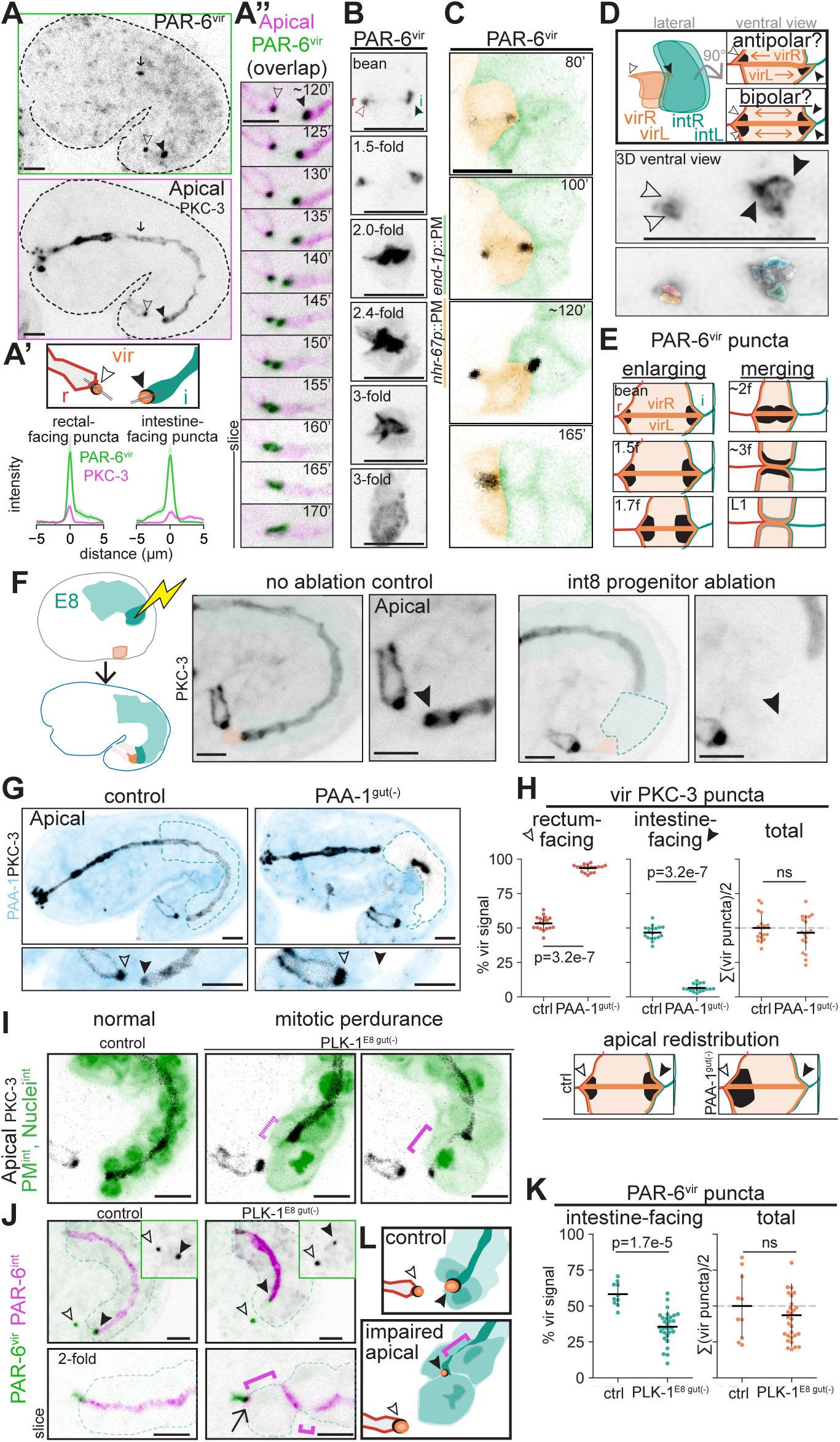
Bipolar rectal valve cells require signaling from the intestine. A) PAR-6^vir^ (black) shows apical surface of 3 FOS-1a+ embryonic cells (one pharyngeal valve (arrow) and vir cells (arrowheads)), colocalizing with rectum-facing (white arrowhead) and intestine-facing (black arrowhead) PKC-3 (magenta) puncta. A’) Line scans (double line in cartoon) of PAR-6^vir^ (green) and PKC-3 (magenta) background-subtracted intensity at both vir puncta (mean and 95%CI of n=12 1.5-fold embryos) A’’), Time course of embryo in A aligned to rectum. Overlap of PAR-6^vir^ (green) and PKC-3 (magenta) signal yields black. Single z-slices upon twitching at 160’. B) 100x images of PAR-6^vir^ (black) in CO_2_-paralyzed embryos. C) PAR-6^vir^ with cells colored by tissue (*end-1p*::PM/*nhr-67p*::PM) and explanatory model (right). D) Optical 168X super-resolution imaging of PAR-6^vir^ substructures with potential models (>2 substructures in 6/6 rectum-facing and 7/8 intestine-facing puncta). E) Model of vir puncta (black) over time (f, -fold). F) Apical signal after posterior intestinal progenitor laser ablation (black arrowhead; vir-intestine contact determined from global *pie-1p*::PM; no ablation: n=10, Eprp/Eplp ablation: n=13). G) Apical surfaces (PKC-3, black) in embryos with intestine-specific depletion of PP2A phosphatase subunit PAA-1 (blue). H) Percentage of total PKC-3 intensity in vir partitioned between apical puncta. Right, total intensity as percentage of mean of control embryos (dashed line). Bottom, cartoon of apical redistribution. I-K) Intestinal mitotic perdurance (*end-1p*::PM, *elt-2p*>H (bipartite transgene, see Methods), green) in control (no degron-tagged allele) and PLK-1^E8^ ^gut^(–) embryos. Apparent gaps in apical (PKC-3, black) signal, magenta brackets. I) Tissue-specific apical signal in PLK-1^E8^ ^gut^(–) embryos. Black arrow, PAR-6^int^ and PAR-6^vir^ signal present at posterior end of intestinal cells. J) PAR-6^vir^ signal quantified as in G. Unless noted, all images are maximum intensity projections of 1.5-fold embryos, quantifications from comma—2-fold embryos, and controls lack degradation transgene. Mean +/-standard deviation. p-values, Mann-WhitneyU. Scale bars, 5 µm.

Following the appearance of the bipolar distribution of polarity proteins, the intestinal and rectal-facing puncta merged into a single apical surface along the virR and virL interface (Figure 3A-C, Figure S4H). Near embryonic hatching and in the L1 larval stage, the PAR-6^vir^ signal became weaker and more unpolarized. Endogenously-tagged PAR proteins were similarly absent from vir in L1 larvae (Figure S4C,D) and junctional proteins localized in a pattern consistent with a dorsal and ventral junction between the two vir cells (Figure S3I). Therefore, the joining of the internal intestinal apical surface and the external-facing rectal lumen occurs by bipolar cells merging apical surfaces along the virR/L interface.

### Intestinal cells are required for valve bipolarity

Given the bipolar localization of the polarity puncta within the vir cells, we hypothesized that these cells gain polarity information from the tissues that they connect. We therefore asked whether the intestine is required for vir cell polarization. Using laser ablation, we removed the precursor to the posterior intestinal cells before the vir cells contacted the posterior intestine (Figure 3F). As expected, the ablated intestinal cells remained as corpses within the embryo but lacked any apical localization of polarity proteins. Strikingly, although cells appeared to contact the intestinal cell corpses, no intestine-facing PKC-3 positive puncta were visible (Figure 3F), indicating that live intestinal cells are required to induce vir polarity at this interface. To further test whether intestinal cells are required for vir polarity, we leveraged a severe phenotype resulting from intestine-specific degradation of the PP2A phosphatase scaffolding subunit^54^ PAA-1 (PAA-1^gut^(–)) in which intestinal cells perdured in mitosis for hours instead of minutes and hatched with an intestinal cyst disengaged from surrounding tissues (Figure S5A-B). PAA-1^gut^(–) embryos were extremely delayed in intestinal polarization (comma – 1.7-fold embryos with disconnected apical patches: control: n=0/18, PAA-1^gut^(–): n=18/18), allowing us to ask whether alive but severely compromised intestinal cells induce vir bipolarity. In PAA-1^gut^(–) embryos, vir cells appeared to contact and push into the posterior intestine yet no intestine-facing puncta were visible and instead all apical signal redistributed to the rectal-facing puncta (Figure 3G,H). PAA-1^gut^(–) intestinal cysts represent a severe phenotype that is a sum of prolonged mitosis and reduced apical polarization, among many other potential effects; therefore, we tested specifically how intestinal cell cycle state (Figure S5C-H), apical polarization, and adhesion (Figure 4, below) affect communication with the neighboring valve as it polarized.

**Figure 4:**
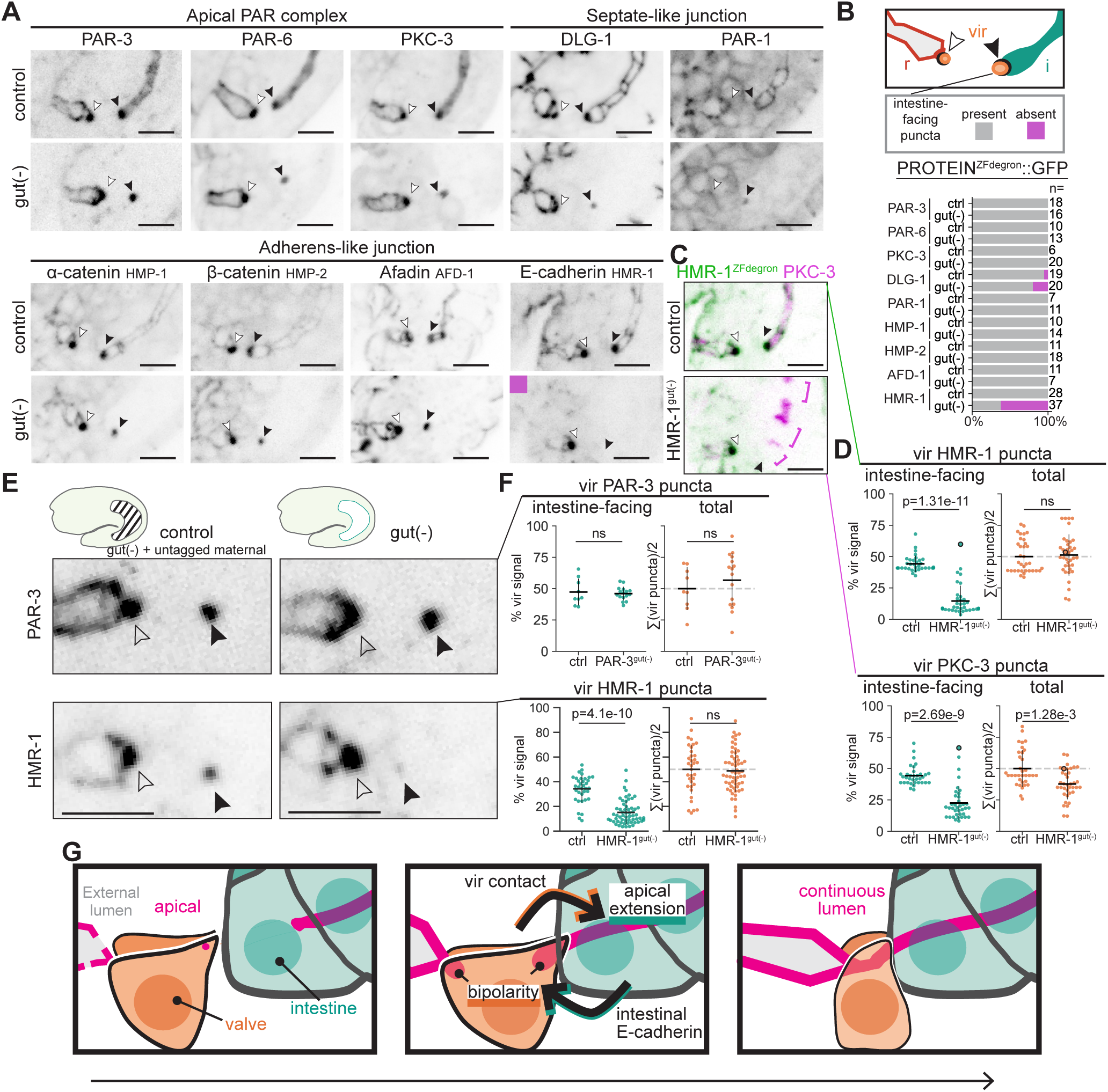
Intestinal E-cadherin is required for valve polarity. A-B) ZF_degron_::GFP-tagged protein localization at the intestine-facing (black arrowhead) and rectum-facing (white arrowhead) puncta in control and indicated gut(-) embryos. C-D) Apical (PKC-3) signal in HMR-1^gut^(–) embryos, with HMR-1 and PKC-3 signal quantified as in Figure 3G. Circled point in HMR-1^gut^(–) graph indicates embryo without intestinal apical gaps. E-F) PAR-3 and HMR-1 intensity with control (gut(-) + untagged maternal) that isolates signal from the intestine-facing puncta, quantified as in Figure 3G. G) Cartoon summary of normal intestinal-rectal connection via reciprocal E-cadherin-mediated signaling between vir and posterior intestinal cells. All images are maximum intensity projections of 1.5-fold embryos. Scale bars, 5 µm.

Since mitosis has previously been shown to be an assault on apical continuity, compounding the defects of losing apical and junctional regulators^55–59^, we asked how intestinal mitoses impact vir bipolarity. Since the valve cells contact the E16 intestine after apical polarization and before the E16-E20 divisions (Figure 1C, Figure S2A-C), we created genetic tools to challenge the intestine to perdure in mitosis during E16 and thereby compromise the apical surface, using intestine-specific degradation of mitotic regulators starting at E8 (‘E8 gut(-)’, *ifb-2p*::ZIF-1^48^). Depletion of mitotic kinases CDK-1, AIR-1/AuroraA, or PLK-1, centrosomal protein SPD-5, or PAA-1 caused variable phenotypes, including fewer intestinal cells, cell rounding, and persistent condensed chromatin (Figure S5D-H). Mitotic perdurance was associated with the appearance of apical gaps in PKC-3 signal, most often in the posterior intestine, that worsened during elongation (Figure 3I, Figure S3F). One of the strongest conditions, PLK-1^E8^ ^gut^(–), had larval lumen obstructions, slowed growth, and delayed intestinal polarization (Figure S5I-L). However, whether the apical gaps in PKC-3 signal were within the intestine or between the intestine and vir was unclear, and so we created a strain labeling the vir (PAR-6^vir^) and intestinal (PAR-6^int^) apical surfaces separately in the PLK-1^E8^ ^gut^(–) background. Of the 18/83 late bean to 3-fold stage PLK-1^E8^ ^gut^(–) embryos examined that had gaps in the posterior apical surface, 14/18 had clear PAR-6^int^ and PAR-6^vir^ signal at the end of the intestine (Figure 3J), indicating that apical gaps developed within the intestinal cells and not between the intestine and vir even in embryos with short and/or discontinuous apical surfaces. In nearly all PLK-1^E8^ ^gut^(–) embryos, PAR-6^vir^ at the intestine-facing puncta was diminished (Figure 3J-K), consistent with an intestine-mediated reinforcement of that apical surface over time. Thus, intestinal mitotic arrest and the associated compromised apical surface attenuated but did not prevent the normal bipolarity of the neighboring vir cells, revealing that the intestine reinforces valve cell polarization over time.

### Intestine-specific E-cadherin is required for valve cell bipolarity

To identify the molecular mechanisms by which the intestinal cells mediate the formation of bipolarity in the neighboring vir cells, we performed a targeted screen of apical and adhesion proteins. Using tissue-specific degradation, we depleted proteins from the embryonic intestine prior to intestinal polarization (gut(-), *elt-2p*::ZIF-1,^25,45,59^) and then examined the intestine-facing vir puncta. Given that our degradable alleles were endogenously tagged with GFP, this screen also allowed us to visualize the endogenous localization at the intestine-facing puncta in the vir cells (Figure 4, black arrowhead) to confirm that the PAR-6^vir^ puncta we observed were not an overexpression artifact. Indeed, all apical and junctional proteins tested localized to intestine-facing puncta within the vir cells that were highlighted against the loss of intestinal signal.

Removal of apical PAR complex proteins from the intestine had no effect on the corresponding intestinal-facing puncta of these proteins in the vir cells, despite severely compromising the intestinal apical surface^25,45,59^. Similarly, degradation of junctional components had little effect (Figure 4A) with the striking exception of E-cadherin/HMR-1. HMR-1^gut^(–) embryos had a strong reduction in both HMR-1 and PKC-3 at the intestine-facing puncta in vir (Figure 4C,D). To quantify the vir portion of endogenous HMR-1 in these puncta, we created a strain in which HMR-1^gut^(–) embryos were rescued by untagged, non-fluorescent, non-degradable maternal HMR-1. This control allowed us to compare endogenous HMR-1 levels in vir at the intestine-facing puncta with and without functional intestinal HMR-1 (Figure 4E-F). The sum of the HMR-1 fluorescence intensity of the bipolar rectal-facing and intestinal-facing vir puncta remained constant with or without functional intestinal HMR-1, consistent with the intestine-facing and rectal-facing puncta drawing from a shared pool of protein components in vir to correctly partition bipolarity. Conversely, PAR-3^gut^(–) embryos in which one of the key determinants of intestinal apical polarity is degraded^20,25^ did not disrupt valve cell bipolarity, pointing to E-cadherin-mediated signaling at the cell-cell contact as the crucial factor coordinating apical alignment across different tissues.

## Discussion

Here we show that cells at the boundaries of connecting epithelial tissues reciprocally exchange orientation information to correctly align their apical surfaces. Using the morphogenesis of the posterior *C. elegans* digestive tract, we defined how a specialized two-cell tissue, the rectal valve, enables the internal intestinal lumen to join the external-facing rectal lumen via E-cadherin-mediated cell-cell interactions (Figure 4G). The process requires two steps: the endodermal intestine transitions from a cyst to a tube and then the exposed intestinal apical surface joins to the ectodermal rectum by merging of bipolar apical surfaces within vir cells.

The reorientation of existing polarity within the posterior intestine involves the extension of apical surfaces along the cell-cell interface to connect with vir cells. This reorientation of existing apical and junctional surfaces where endodermal tissue abuts ectodermal tissue, such as those observed in pharyngeal arches^60^ and foregut^61^, is a widespread feature of development that is not well understood. Our data point to separable pathways for polarity establishment and tissue connections, as PAR-3^gut^(–) embryos displayed normal vir bipolarity, despite drastic apical polarity defects within the intestine^37^. Connections at tissue boundaries in systems from mammalian kidney to *Drosophila* testes similarly appear to be under separate genetic control than the initial process of apical polarization^46,62–65^. However, future studies using tissue-specific approaches will be required to identify general principles of how polarity establishment interfaces with pathways controlling tissue connection.

Vir cells establish bipolarity as they integrate with the already polarized intestine, transitioning along the mesenchymal-epithelial continuum. This integration is similar to radial intercalation of unpolarized cells into existing epithelial tubes and sheets^66–69^ except that vir cells experience a competing source of apical orientation information from contact with rectal cells. Cells at tissue-tissue boundaries often integrate competing sources of polarity information, whether they lose polarity and then repolarize as in kidney and optic fissure closure^38,70^ (analogous to early vir polarization upon intestinal contact) or they maintain aspects of polarity while simultaneously invading into the neighboring tissue as in avian lung, mouse axial mesoderm, and ascidian branchial fissures^60,66,71^ (analogous to later bipolar vir cells). Even lumen formation in single primordia can involve the fusion of multiple cysts, and polarizing cells at the edges between cysts find themselves in a bipolar environment before apical alignment is complete^72–76^. Vir cells polarize with input from the intestine and rectum via E-cadherin signaling, which we found to be independent of its role in catenin-mediated adhesion^45^. Thus, vir cells are a model for the formation of this understudied bipolar state during tissue connection.

We speculate that developmentally appropriate bipolarity, putatively reflecting a partial mesenchymal-partial epithelial state, may be hijacked by carcinomas to drive metastatic cell behaviors, as cell clusters and cells with partial epithelial-mesenchymal transitions are the most likely drivers of productive metastases^77–79^. Additionally, E-cadherin is present on invadopodia^80,81^, suggesting that signaling at cell-cell contacts and the transfer of apical orientation information is important in not only developmental but also disease contexts.

## Acknowledgements

We would like to thank John Murray for alerting us to *fos-1* specificity, Michael Nonet and Callista Yee for sharing reagents prepublication, Abby Converse for cloning, Ben Snyder for code refactoring, CGC for strains, Nikon for super-resolution microscope demonstration, Victor Naturale for scientific input, and Feldman lab members for feedback. LEC is a Damon Runyon Fellow supported by DRG-2428-21; support from K99 GM13548901 to MDS, NSF predoctoral fellowship and T32 GM007276 for RKN, F32 GM129900-01 to MAP, R01GM136902, R01GM133950, R35GM153310 to JLF.

## Author Contributions

Conceptualization, J.L.F. and L.E.C.; methodology, L.E.C., M.D.S., J.M., J.L.F.; formal analysis, L.E.C., M.D.S.; investigation, L.E.C., J.L.F., M.D.S., R.K.N., M.A.P.; writing— original draft, L.E.C. and J.L.F.; writing—review and editing, L.E.C., J.L.F., M.D.S., R.K.N., M.A.P.; visualization, L.E.C.; supervision, J.L.F.; funding acquisition, J.L.F., L.E.C., M.D.S., R.K.N., M.A.P.

## Supplemental Figure Legends

**Figure S1:** Rectal valve (vir) cells facilitate the connection between the intestine and rectum. A-C) Maximum intensity projection confocal images of live 1.5-fold stage embryos with single channel insets at right, outline corresponds to channel. A) Image of digestive tract membranes (*pha-4p*::GFP::CAAX(PM), green) and apical surfaces (endogenously tagged PKC-3::RFP, magenta), unspliced version of image shown in Figure 1A. Right, tissue annotation (p: pharynx, i: intestine, v: valve, r: rectum). B) Intestinal cell membrane (*end-1p*::GFP::CAAX(PM), green), subset of pharyngeal and rectal cells (*ceh-27p*::LifeAct::mScarlet, magenta), and brightfield (BF) image show location of these tissues with single channel insets at right. The indentation in the posterior-most intestinal cells (int9) was visible in 14/14 embryos, while single-copy *ceh-27p*::LifeAct transgene (predicted expression^50^ in K/K’, PVT/rectD, virL/R and their progenitors) was visible in the valve cells only in 6/14 embryos. C) Apical surfaces (PKC-3::RFP, magenta) surround an externally secreted apical extracellular membrane component^82^ (LET-653::GFP, green) at the developing mouth/buccal cavity and rectum but not the internal pharyngeal or intestinal lumens. n=13. D) The two rectal valve cells (*nhr-67p*++ magenta, FOS-1A+ green, co-expressed in 38/38 embryos) lie on top of each other in 1.5-fold embryos. A single slice from a 100X image at two different z planes show virL and virR relative to basement membrane (LAM-2/Laminin::mNeonGreen green). M, intestinal midline where apical proteins localize. Initially basolateral proteins also localize at the midline^25^, including secreted Laminin. E) Confocal image maximum projections through each valve cell (2.5µm and 1.5µm respectively) at 1.5-fold stage. Co-expression of strong *nhr-67p*::H1::mCherry signal and *pha-4p*::PM, n=42/42 comma-1.7-fold embryos. Arrows point to membrane extension of valve cells into the indentation between int9R and int9L. F) Longitudinal sections from electron microscopy of a 16 hr L1 larvae^27^ with 0µm set to slice 868, which shows the valve cell lumen. Cartoon at top modified from wormatlas.org to show cells of larval intestine. G) Rectal length in L1 larvae after indicated ablation, colored by intestinal lumen phenotype. ns, not significant one-way ANOVA. Example L1s imaged 1 day after indicated laser cell ablation of progenitor cells in the embryo. Right-most vir progenitor ablation is full overview of larvae shown in Figure 1D. Left-most rectD progenitor ablation example is undergoing expulsion in the defecation cycle, showing the intestinal and rectal lumens fully open. Single channel underneath shows apical PKC-3::GFP (magenta) localization. Unless noted, all images are maximum intensity projections of 1.5-fold embryos, quantifications from 1.4-1.6-fold embryos. Scale bars, 5 µm.

**Figure S2:** The rectal valve vir cells undergo morphogenesis to integrate with intestine. A) Detailed timeline of embryonic development and expression of markers within the valve cells (bottom). Legend shared between G and H. B) Cell-cell contact area (y-axis) plotted over time (every ∼1.5’) from whole embryo segmentations^36^. Light orange, contact of posterior-most valve cell with intestine (sum of ABprpapppp(p) and Epxpp(a)). Dark orange, contact area of anterior-most valve cell (sum of ABprpappppa and Epxpp(a)). The time of cell divisions (triangles) and other morphological events are shown for the 4 embryos that show bilateral virR/L cells. Timepoints with clear segmentation errors are not plotted. mpf, minutes post fertilization. C-G) Examples of morphogenetic events from live confocal imaging (intestine: C-E, valve: F-G) with cells labeled as in Legend. Time 0’ is set to vir progenitor division into virR/virL. C) Lateral view of intestine with apical marker PKC-3::RFP at the intestinal midline (M). D) Lateral view of intestine undergoing int5 intercalation (same embryo as Figure 1F). E) Lateral view of intestine undergoing posterior ‘star cell’ division to create int9. F) Two single optical sections show cells on and 6.5µm below the ventral surface undergo gastrulation (ABprpapppa (unmarked), ABprpapppp (orange, valve progenitor), ABpxpppppa (grey open circles, muscle)) during early E16 oriented with anterior up, ventral view. Gastrulation end is defined as contact between ABplpapppp (shaded red, PVT/rectD progenitor) and ABprppppaa (shaded red, PVP/rectR progenitor). G) The anterior-most valve cell virR rotates from anterior to posterior so that the valve cells are bilaterally symmetric at the midline of int8 cells, as shown by two separate examples of vir markers that also mark the intestine(*pha-4p*::PM or NCAM-1::GFP). Both valve cells become characteristically wedge-shaped and remain wedge-shaped throughout the rest of bean stage. H) Valve cells do not breach a basement membrane (representative of n=28 time courses across 4 strains). Ventral view of 3D rendering of LAM-2::mNeonGreen shown with outlines of vir cells drawn from *nhr-67p*::PM outlines of GFP::FOS-1A+ cells. Box indicates location of detailed ventral view time course on right and insets in L,M. I-K) Maximum intensity projections of 1.5-fold embryos with junctional, basolateral, and apical markers in the digestive tract. I) vir cells have the junctional/basolateral Scribble-homolog LET-413 on all visible membranes in 11/11 embryos. J,K) There is a gap in both apical and junctional signal in between the two large puncta (n = 45/45 1.5-fold embryos) that fills in during elongation (n = 8/8 time courses). Timecourse is aligned to rectum. Unless noted, scale bars, 5 µm.

**Figure S3:** The intestinal cyst-to-tube transition. A-C) Cyst-to-tube visualizations. 0’ set to approximate completion of cyst-to-tube transition. A) 3D rendering of apical PKC-3 (black) with 3D cell shapes manually segmented from *pha-4p*::PM (green, bottom). B) Example sparse timecourse of single embryo undergoing cyst to tube transition with 3D rendering of lateral view and two 0.5μm optical z-slices from locations shown ventral view. Double lines show location of int8L/R interface and arrow marks intestinal apical surface. (n = 16/16 embryos completed cyst to tube transition before pharynx polarized into a straight line (E16 stage 3^25^). C) Example time course of cyst to tube transition (30-35’) in embryo expressing HMR-1::mScarlet (green) and PAR-3::GFP (magenta) shown as maximum projection of resliced image through apical surface (n=6/6 embryos completed cyst to tube transition before E16 stage 3). D) Laser cell ablations of vir progenitor and control neighboring cell (top: pre-ablation, red and post-ablation, blue.) Arrow points to targeted cell. Note debris raised in DIC and not loss of fluorescence was used to confirm cell killing. Those embryos at comma stage appeared to have the PKC-3+ apical surface stop before the end of the intestine (bottom, control ablation n=6, vir ablation n=10). Larvae resulting from this experiment included in Figure 1D, S1N and microvilli marker is not expressed strongly until 2-fold^83^. E) Additional lateral single slices of embryos after laser cell ablation shown in Figure 2C. F) *nhr-67p*::H1 is expressed in cells that line the ventral cleft (red, ventral surface; blue, 1.5μm below surface). White asterisk marks vir progenitor ABprpapppp. G) Validation of degradation extrachromosomal array wowEx128 using degradation of SPD-5::ZF::GFP^84^ to block mitosis in nhr-67+ cells (ventral closure defects (arrow): control n=0/5, SPD-5^nhr-67p^(–) n=12/13). H) Additional examples of HMR-1^nhr-67p^(–) intestinal phenotypes. I) HMR-1 intensity and BFP::CAAX intensity (expressed from *nhr-67p*::ZIF-1:: HMR-1::E2A::BFP::CAAX) were measured at vir lateral membrane (grey circles) and colored based on phenotype (colored bars in Figure 2E,F and Figure S3H) combined from 4 independent experiments. Unless noted, all images are maximum intensity projections of 1.5-fold embryos, quantifications from comma-1.6-fold embryos. Scale bars, 5 µm.

**Figure S4:** vir cells are bipolar. A) A valve-specific promoter was found by examining the expression patterns of GFP-tagged *fos-1* fosmid, *fos-1* C-terminal endogenous GFP tag, and *fos-1a* N-terminal endogenous GFP tag. B) vir-specific apical reporter (PAR-6^vir^) was created by inserting a PAR-6::mNeonGreen::P2A transgene at the start codon of the endogenous *fos-1a* locus to create allele wow181. Embryo is overview of 168X super-resolution image in Figure 3D showing the apical surface of 3 embryonic cells (one pharyngeal valve cell, arrow and the two rectal valve cells, arrowheads) Note the non-specific gut-granule autofluorescence in the intestine (green outline). C-E) PAR-6^vir^ (black) localization throughout development, oriented with rectum to left and intestine to right. C) Categorization of PAR-6^vir^ localization patterns and embryo stage (n = 215 embryos imaged across 5 strains). D) Example live images from CO_2_-paralyzed embryos or levamisole-paralyzed L1 larvae with colored square corresponding to localization categorization in C. Scale bars, 2μm E) Three representative time courses of PAR-6^vir^ puncta appearance (n = 8) and merging (n = 13). All images, unless noted, in ventral view to see interface of both vir cells (virR on top). Colored bars at right correspond to localization categorization. Time course on left shows unspliced version of Figure 3C. Scale bars, 2μm F) Labeled cells in bean stage serial section EM dataset (320 min post first cell division (mpfc) XY section (541,702,257)) and comma stage (345mpfc YZ section (1152,807,199)) 65nm sections^85^. Arrows point to contralateral contacts between vir and int cells. G) Labeled cells in transverse 50nm sections from ∼550 min 3-fold embryo^53^ (RDE sections #25, #31) sections. Arrows point to contralateral contacts between vir and int cells. H) Example live CO_2_-paralyzed 3-fold stage embryo with junctions outlined by DLG-1 (cartoon of junctions in rectal (red), vir (orange), and intestinal (teal) cells on right) and PAR-6^vir^ localized along L/R interface. I) In L1 larvae, apical signal from endogenously-tagged PAR-6 is not apparent in vir cells (n = 13/13 larvae). Endogenously-tagged PKC-3::RFP (magenta) in vir shows no signal (8/17 larvae) or a single pixel of signal (9/17, arrow) while endogenously-tagged HMR-1::ZF::GFP (green, appears black in merge) along with PKC-3 is visible as a dorsal and a ventral line presumably along virL/R interface. Intestinal cytoplasmic GFP (*ges-1p*::GFP) helps visualize the end of the intestinal cells. Unless noted, all images are maximum intensity projections of 1.5-fold embryos. Scale bars, 5 µm.

**Figure S5:** Consequences of mitotic perdurance in the intestine. A-B) Fixed embryos immunostained using antibodies to highlight mitotic cells (11pH3, magenta) and live control and PAA-1^gut^(–) (PAA-1::ZFdegron::GFP; *elt-2p*::ZIF-1^degrader^::sl2::mCherry::H2b) larvae. Control embryos are the same genotype as PAA-1^gut^(–) except the *paa-1:zf:gfp* locus is balanced by hT2. A) pH3+ cells in mid-intestine (control: n = 0/20; PAA-1^gut^(–): n=23/23 bean – 1.7-fold stage embryos). PAA-1 channel (green) LUT individually adjusted to show embryo morphology. B) Example larvae with intestinal cyst: (control, n=0/14 PAA-1^gut^(–), n=9/9 larvae). Blue arrow marks end of pharyngeal lumen and red arrow marks end of rectal lumen. C) Cartoon summarizing effect of persistent mitotic arrest on the intestinal apical surface. D) Cartoon depicting intestinal cells dividing from E16 to E20 (stars). E) Legend for F (colors only) and G (colors and shapes). In normal development, int1 and int8 cells briefly enter mitosis and have continuous PKC-3 throughout the mitotic process^34,59^. Embryos were called as having “mitotic” cells if at least one cell in int1 or int8 pair showed condensed chromatin and/or had a rounded cell shape. After E20, in the final form of the intestine, int1 ring is formed from 4 cells while int9 is formed from 2 cells and abnormal larvae (magenta) were called as having ‘1-cell’ if the posterior-most part of the intestine had intestinal PM signal containing 1 nucleus whereas ‘2-cell’ posterior-most intestines had a clear juxtaposition of two cells forming a ring. F) Categorization of embryos (left) and larvae (right) with n to right of the graphs. Unless noted, control in Figure S5F-L is a strain with indicated markers and *ifb-2p*::ZIF-1^degrader^ without a degron-tagged allele. Embryos were evaluated at both the most anterior cell pair (“A* int1L/R”) and most posterior cell pair (“P* int9L/R”) for mitotic cells and apical gaps in PKC-3 signal in strains with and without a transgene labeling intestinal chromatin (‘*elt-2p*>H’, recombined elt-2::tetR-QFactivating domain, tetO::GFP::histone)). Larvae were evaluated at the most posterior end (“int9”) for cell number (*end-1p*::GFP::CAAX(PM) and *elt-2p*>H required for most accurate cell counting) and apical gaps in PKC-3 signal. Note that PAA-1^E8^ ^gut^(–) larvae were only scored for continuous vs apical gap as the strain used for imaging lacked *end-1p*::PM (hatched shading). G) Total intestinal cell number over developmental stages for each indicated genotype shown as mean +/-s.d. for each condition. All datapoints are from animals with both *end-1p*::PM, *elt-2p*>H, except for PAA-1^E8^ ^gut^(–) condition which includes some embryos and all larvae with only *elt-2p*>H. The datapoint for each embryo is colored/shaped by the phenotype of its P* cells as in legend Figure S5E (for embryos: “P* int9L/R” apical gaps and mitotic state and for larvae: “int9” apical gaps and cell number). n embryos assayed listed on bottom of graph. Note that beeswarm-style resulted in suppression of plotting up to 1/3 of individual points at modal values (all non-mitotic, no gaps). H) Example embryos expressing apical (PKC-3, black) and intestinal markers (*end-1p*::PM, *elt-2p>*H, green, LUT adjusted individually to visualize intestinal morphology). Apical gaps are indicated by magenta brackets and mitotic cells by grey circles. I) Lumen continuity assayed by feeding L1 larvae dyed food. Phenotypic categories quantified on left. Example larvae at right (arrows point to anterior portion of rectum that fills with food when lumen is continuous). Controls are the degradable allele alone without overexpression of the degrader transgene. J) Larval growth at day 3 for indicated genotypes. ‘control’ lacks both degron-tagged alleles and degrader transgene. AIR-1/PLK-1 control as in Figure S5I. K) Example larvae showing apical gap (magenta bracket) in PKC-3 (black) and 1-celled int9 ring. Asterisk marks cyst. Intestinal markers show both nuclei and membrane (green, gamma=0.25). L) Example sparse time-courses showing apical signal throughout E16-E20 in individual embryos (embryos with nearly-continuous midline-associated PKC-3 (E16 stage 2.5^25^) with pharynx beginning to polarize (0’) – control: n=14/14. PLK-1^E8^ ^gut^(–): n=2/23). Of the 21 PLK-1^E8^ ^gut^(–) embryos with delayed polarization, 15 formed gaps in PKC-3 signal (magenta brackets) at the posterior with 7/15 gaps clearly within the intestine and the rest unclear. Unless noted, all images are maximum intensity projections of 1.5-fold embryos. Scale bars, 5 µm.

## Supplemental Video Legends

Video S1: Segmentations from Guan et al, 2025 over time from ventral and lateral view showing the lineage of vir (orange) and posterior intestine (yellow) within the context of the entire embryo through comma stage.

Video S2: Puncta merging from 1.5 – 2-fold stage embryo from Figure 1H, showing the apical highlight of PKC-3 signal.

Video S3: Movie of the cyst to tube transition in pre-bean/bean stage embryo from Figure 2A. Bottom views show 3D renderings of cells and/or interface at every other time point.

## Methods

### Resource availability

#### Lead contact

Further information and requests for resources and reagents should be directed to and will be fulfilled by the lead contact, Dr. Jessica L. Feldman.

#### Materials availability

Code and source data for analysis, plasmid maps, example raw image z-stacks and processed images for figures available at doi: 10.17632/7n37yyb9vz.1 . DNA constructs and transgenic *C. elegans* strains generated in this study are available from the Lead Contact, Dr. Jessica Feldman, upon request.

### Experimental model and study participant details

#### Husbandry of experimental animals

*C. elegans* strains were cultured on Nematode Growth Medium (NGM) plates coated with a lawn of E. coli OP50^86^ unless otherwise stated. For all experiments, animals were maintained at 20°C. Strains used in this study are listed in the key resources table. For all experiments, embryos were obtained from hermaphrodites. Embryos were either harvested from the uterus of young day 1-2, gravid adult hermaphrodites that were incubated in M9 media or were picked directly from culture plates after they were laid by hermaphrodites. For strain generation that required genotypic mixing, male and hermaphrodite strains were crossed. 5 adult male worms and 3 L4 hermaphrodite were incubated on 3.5 cm NGM plates seeded with OP50 and cross-progeny were isolated and the desired genotypes were identified either through direct visualization of markers when possible or by PCR genotyping.

### Method details

#### vir marker identification

Databases of expression in lineaged *C. elegans* embryos^50,51^ were mined for expression in ABprpapppp and marker discovery between ABprpapppp and ABplpapppp from lineage-labeled single-cell sequencing data^52^ followed by visual confirmation in embryos yielded *nhr-67p* (stIs10684), *ceh-27p* (stIs10440), *pha-4p*::PM (pxIs10), NCAM-1 (wow107), FOS-1A (bmd138) as markers of vir cells.

#### Transgenes and endogenous alleles

Cloning strategy for each plasmid, primers, and injection mixes for each transgene/allele detailed in Supplemental Table 1. All endogenously-tagged alleles were created using the self-excising cassette (SEC) CRISPR/Cas9^87^ method except wow185 DLG-1::mScarlet-I, created via ribonucleoprotein CRISPR/Cas9^88^. Cloning was performed using Gibson Assembly (NEBuilder HiFi DNA Assembly Master Mix, NEB) using digested backbone and genomic PCR products amplified with touchdown PCR or geneblocks (IDT). The ZF::GFP SEC repair template pJF250^48^ was modified to create additional ZF::FP backbones (pLC001 ZF::GFP11, pLC015 ZF::BFP, pLC019 mScarlet::FLAG) and grown in ccdB survival stain (Invitrogen). At least six individual minipreps digested with appropriate enzymes (N terminal tags SpeI/ClaI ; C terminal tags AvrII/SpeI) were verified on an 1% agarose gel to not have plasmid recombination before combining for use in HiFi reaction with 0.025 pmol of each component for 4 hours 45°C to create gene-specific repair templates with 0.5-1kb homology arms. For sgRNA plasmids and pLC150[*end-1p*::PH::mCherry], site-directed mutagenesis (Q5 SDM kit, NEB) was used to generate plasmids. For SEC-based CRISPR edits, Cas9 and sgRNA were delivered by co-injection with gene-specific plasmid^87,89^, isolated by following rol phenotype before heat-shock to remove SEC, and backcrossed 2x to JLF155 *zif-1(gk117)* III. To create PAR-6^vir^ transgene, the ‘ZF::BFP’ of the SEC repair template for ZF::BFP^SEC^::FLAG::FOS-1a was replaced with ‘PAR-6::mNeonGreen::P2A’ and injected with published sgRNA^90^ so that the fos-1a locus produces separate PAR-6::mNeonGreen and FLAG::FOS-1 proteins after riboskipping. Plasmids to create single-copy transgenes wowSi15 (*elt-2p*::tetR-AD) and wowSi31 (*ceh-27p*::LifeAct::mScarlet) were made with GoldenGate assembly using SapI or BsaI (NEB) ligation^91,92^. Single-copy transgenes were made using FLP recombination-mediated cassette exchange^91^ and the bipartite transgene (‘*elt-2p*>H’ wowSi41) made with phiC recombination^92^ of elt-2::tetR-AD and tetO-driven marker of condensed chromatin. Gene diagram of *fos-1* locus in Figure S3B from http://www.wormweb.org/exonintron.

#### Tissue-specific degradation alleles and controls

All degradation strains were maintained on a balancer (with the exception of AIR-1^E8^ ^gut^(–)) and all degradation are maternal/zygotic degradation (embryos imaged are from balancer negative mothers). For strains readily viable off of a balancer (all strains except PAA-1gut(-), PAR complex gut(-), and HMR-1gut(-))^25,45,48,59,84^, some balancer negative mothers of imaged embryos were 1-2 generations off the balancer.

Degradation transgenes to generate gut(-) animals have all been previously validated (see strain list). Degradation in *nhr-67p*+ cells (neuroblasts, rectal cells) via wowEx128 was verified using SPD-5::ZF::GFP and quantification of HMR-1 degradation. Note that transmembrane proteins such as HMR-1::ZF::GFP seem to require higher expression of ZIF-1 to yield strong degradation than other ZF::GFP proteins. Controls for degradation are the same strain lacking the degradation overexpression except for analysis of strains with *end-1p*::PM and/or *elt-2p*>H in which ‘control’ is degradation overexpression without the degradable allele. Maternal untagged control in Figure S3J and Figure 4E are hT2-embryos from hT2+ mothers who provide untagged protein. hT2 is a GFP-marked translocation of chromosome I and III, so GFP-embryos are ¼ euploid embryos with only maternal untagged protein that are able to survive to adulthood and ¾ aneuploid embryos with maternal untagged protein plus zygotic untagged protein from GFP-portion of the translocation with uncharacterized lethality.

#### Antibody staining

Embryos dissected from one-day old adults of desired genotype incubated in M9 were attached to microscope slides coated with poly-lysine and containing Teflon spacers. Slides were frozen on dry ice, embryos were permeabilized by freeze-crack method^4^ and submerged in 100% MeOH for 5-10 minutes at −20°C. Rehydrated embryos were incubated in 1:500 anti-pH3 (Ser10) overnight at 4°C, washed three times in PBT, incubated in secondary antibody (anti-rb-Alexa-647 1:500) and 500ng/μl Hoescht 33342 for 1 hour at 37°C, washed once in PBT and twice in PBS before mounting in Vectashield (Vector Laboratories).

#### Embryo and larval imaging

Embryos for microscopy were dissected from gravid hermaphrodites incubated in M9 or collected from unstarved plates containing 1-2-day-old gravid adults. For live imaging, embryo samples were mounted on a pad made of 3% agarose dissolved in M9 under a #1.5 coverslip. Live and fixed imaging was performed on a Nikon Ti-E inverted microscope (Nikon Instruments, Melville, NY) using a 60× Oil Plan Apochromat (NA = 1.4) or 100× Oil Plan Apochromat (NA = 1.45) objective and controlled by NIS Elements software (Nikon). Images were acquired with an Andor Ixon Ultra back thinned EM-CCD camera using 405 nm, 488 nm, or 561 nm imaging lasers and a Yokogawa X1 confocal spinning disk head equipped with a 1.5× magnifying lens. For live imaging, images were taken at a z-sampling rate of 0.5 μm for experiments looking at 1.5-fold embryos and 0.3 μm for experiments targeting bean-stage embryos. For fluorescence imaging of L1 larvae in Figure S3S, L1 were picked into a drop of 2 mM levamisole on a 5% pad to minimize movement. Images were processed the Fiji distribution of ImageJ (“Fiji”) v2.16.0/1.54p^93^ or napari v0.4.19.

#### Laser cell ablations

A Micropoint dye laser (coumarin 435 nm) was targeted at desired cell nucleus (20-50x 50-55% power until debris was raised in DIC) in ventral-mounted embryo. Desired *ceh-27p*+ or *nhr-67p*+ cells were identified by location and cell ablation was confirmed by 1) presence of extra-embryonic nuclei, 2) loss of 2 *ceh-27*+ cells at 1.5-fold, or 3) loss of *nhr-67p*++/*pha-4p*+ wedge-shaped cells at late bean stage. Note that ablation of ABprpapppp before a cell-contact mediated inhibition of vir cell fate in ABplpapppp can result in ABplpapppp transforming into vir cells^94^. To facilitate tracking of ablated embryos, all non-ablated embryos on the pad were exploded with high power micropoint laser and embryos were left on pad overnight at 20°C in a humid chamber. Larvae were imaged as above except without the addition of a paralytic agent before rescuing to a OP50-seeded plate. Rectal length for each larvae was averaged from 3 measurements from single z-slices since larvae were not paralyzed. Larvae one day after laser cell ablation were blind scored as ‘continuous, expanded, or cyst’ and the continuous and expanded categories were combined as ‘connected’ for Figure 1D (data combined from following independent experiments - no ablation: 2 experiments,;ABplpapppp: 1 experiment (examination by EM after PVT progenitor ablation^53^ showed that the K.K’ cells moved forward to connect to virL/virR in place of rectD); ABprpapppp: 3 experiments)

#### Imaging chamber for CO_2_ immobilization

The imaging chamber for CO_2_ immobilization was designed using AUTODESK Tinkercad (https://www.tinkercad.com/) to improve upon previous embryo CO2 immobilization setups^95^. The 3D printing files (.stl) are available at NIH 3D 3DPX-022973. The chamber comprises two parts: a bottom part with a square chamber that transforms into a bottom coverslip chamber connected to an exit and entry tubing, surrounded by a moat to house an O-ring; and a top part that seals the chamber’s O-ring. The device was printed on a Digital Light Printing 3D printer Photon Ultra (Anycubic, Inc.) using a UV curing stereolithography (SLA) resin. The printing process involved 3 seconds of on and 1 second of off exposure with a z-step of 50 µm and an XY pixel size of 80 µm. After printing, the two parts were cleaned with 100% isopropanol, and a final cure was performed using a UV blue-light LED for 15 minutes. The device was then dried in an oven for 30 minutes. The top and bottom imaging windows were added and sealed using UV clear resin and 22×40mm #1.5 coverslips. A nitrile rubber O-ring (32×28×2 mm) was added to the chamber’s moat to create a seal. Both the top and bottom parts were lined up with N52 5×2 mm round magnets. Embryos were applied to the coverslip in M9 buffer and overlaid with 3% agarose pad. For immobilization, CO_2_ was allowed to flow through an ozone water humidifier and into the chamber for 5-10 minutes at a rate of 60-80 cc/min until immobilization. The chamber was then sealed by pinching the tubing at the entry and exit valve. After the imaging was complete, ambient air was inserted into the chamber using an aquarium air pump through a flow derivation system.

#### SoRa super-resolution imaging

Optical super-resolution imaging was performed on a Nikon CSU-W1 spinning disk confocal microscope with a 60X 1.42NA oil lens and 2.8X SoRa microlens emission disk. Richardson-Lucy 3D deconvolution using 10 iterations to yield an image with voxel sizes of 0.04um x 0.04um x 0.2um. Images were manually segmented and visualized in 3D using iso_catagorical rendering.

#### Larval assays

For larval growth assay, 10 young balancer negative mothers of desired genotype were placed on small plates for 4 hrs at 20°C in triplicate to lay eggs on day 0. On days 2-4, larvae were counted and L4s/adults were counted and removed. For Smurf assay, L1 larvae were incubated for 3-5 hours in a humid chamber in 30μl OP50 + 10μl 20% blue dye before mounting and scoring by eye. For example larvae for the ‘Smurf’ feeding assay in Figure S3Q, a Vankey Cellphone Telescope Adapter Mount (Amazon) was used with an Apple iPhone 7 or 12mini camera and the compound microscope 60× objective. Brightness, contrast, and hue were adjusted for each color image with Adobe Photoshop v21.0.1.

#### Phenotypic categorization

Number of 1.5-fold embryos with gap between rectum and intestine was determined from maximum intensity projections. Images were visually inspected for arcade cell connecting at the buccal cavity and clear gap between the intestine and rectum indicating that this apical surface is the final surface to become continuous. Embryos examined from all ‘wild-type’ and control conditions containing following endogenously-tagged alleles of PKC-3 (wow118, it309, wow85, cp41), PAR-3 (wow59, it300), HMR-1 (wy1254, wow193), DLG-1 (wow185, wow117). For mitotic regulator characterization: ‘Apical gaps’ in embryos were scored visually by looking for absence of signal along the midline, as determined from plasma membrane signal, of the anterior-most or posterior-most cells. The plasma membrane marker zuIs70 is associated with 21-cell intestines (19% of control L1) due to extra division in the mid-intestine after E20^29,96,97^. ‘Mitosis’ was scored when at least one of the L/R pair of cells appeared rounded and/or had condensed chromatin. ‘1-cell’ or ‘2-cell’ intestinal ring was scored using plasma membrane marker surrounding the most posterior intestinal nuclei. For Figure 4B: intestine-facing puncta presence/absence was scored from maximum intensity projections through the intestine with auto-adjusted brightness/contrast (saturated=0.35). DLG-1gut(-) numbers are the sum to two separate degradation lines: intDeg and wow

#### Localization patterns

Localization patterns for endogenously-tagged proteins in non-disrupted genetic backgrounds were examined in >30 embryos.

Figure S2H Laminin timecourses second marker: *nhr-67p*::PM (JLF1402) n=6, PKC-3 (JLF1469) n=7, *ceh-27p*::LifeAct (JLF1391) n=9, NCAM-1 (JLF1378) n=6.

Figure S4C PAR-6^vir^ localization quantification is from PAR-6^vir^ single images: RFP::PKC-3 (JLF1440) n= 70 immobilized with CO_2_, DLG-1::mScarlet-I (JLF1424) n=64 immobilized with CO_2_, mScarlet-I::CAP-1 (JLF1798) n=38, immobilized due to high concentration of bacteria, PAR-6^int^ (JLF1776) n = 29, imaged before twitching, *nhr-67p*::PM/*end-1p*::PM (JLF1426) n=18 no immobilization. Observation of PAR-6^vir^ puncta appearance in 8 timecourses (PKC-3 n=4*, nhr-67p*::PM n=4) and puncta merging in 13 timecourses (PKC-3 n=2, CAP-1 n=7, *nhr-67p*::PM n=4).

#### Visualization

‘Slice’ corresponds to single 0.3 or 0.5um optical imaging plane and was chosen to best visualize the cells/structures of interest. ‘Max’: Maximum intensity projections were taken through the entirety of the tissue of interest for visualizations. ‘3D’: 3D projections are screenshots of napari 3D rendering using mip (min/max intensity projection of inverted/greyscale images) with camera angle kept constant across time. 3D outlines of cell boundaries/interface were drawn manually on each z-stack before rendering as ‘iso-categorical’ for entire cells (Figure 2A, Figure S3A) and ‘translucent’ to flatten the 3D model for laminin timecourse (Figure S2H) and interface in Figure 2A. For all timecourses, unless noted, 0’ is vir cell division (or within 5’ as estimated from tail spike progenitor divisions) and 1.5-fold stage is set to ∼120’ post vir cell division. Videos were created from Cellular Morphology of *C. elegans* Embryo database^36^, napari-animation, Fiji distribution of imageJ, and annotated in Adobe Photoshop before final rendering. Imaging parameters and linear contrast adjustments were consistent between images compared across time and/or genotype and were applied to entire image with the following exceptions:

Figure 1A and Figure 3C colored by tissue: Multiple versions of PKC-3/PAR-6^vir^ (black) with colored PM channel (cyan-I, Forest-I, BOP orange-I) channel were cropped to desired tissue in Adobe InDesign and spliced together. Unspliced single-channel images are shown in Figure S1A and Figure S4E.

Figure 1H, Video S2 : Application of ‘highlight’ to the image shown in Figure 1G. Intensities within a mask manually drawn along the apical surface were adjusted to a suitable level and added to original image in which nuclei were not oversaturated. This combined image was then rendered in 3D over time to visualize the merging of the apical surfaces along with the rectal nuclei.

Figure S1C: Median filter (despeckle) applied to Figure S1C due to low expression of LET-656.

Figure S2K: Rolling-ball background subtraction (250 pixel radius) was applied to remove uneven background outside of the eggshell before contrast adjustments in order to show HMR-1/PKC-3 puncta merging.

Figure 2A,B, S3A, Video S3: See ‘cyst-to-tube’ transition below.

Figure S4H: Mitotic regulators example 1.5-fold embryos have the green intestinal end-1p::PM, *elt-2p*>H channel adjusted individually to enable visualization of intestinal morphology in each image. PKC-3 contrast adjustments were applied uniformly.

In all images shown and in all quantifications, gamma=1.0 except as listed in figure legends in cases where both bright nuclei and dim apical/PM needed to be visualized in the same channel: vir ablation 3D projections (Figure 2C: JLF914 *nhr-67p*::H1 and PKC-3, gamma=0.5 magenta LUT imaged in 561nm channel,) and PLK-1^E8^ ^gut^(–) larvae (Figure 3S: JLF1527 and JLF560 end-1p::PM, *elt-2p*>H, gamma=0.25 forest-I LUT imaged in 488 channel).

### Data analysis

#### Timeline construction and cell-cell contact quantification

Cell-cell contact was plotted from segmentation data^36^ using a min post fertilization (mpf) offset to 4-cell stage from which all 8 segmented embryos were originally aligned in time (estimated to be 110’). Cell-cell contact between int8R and int8L examined to identify timepoints with segmentation errors that were removed from graphs. Division timepoints and cell-cell contact area data were taken from published repository and contact between desired pairs of data was summed. ‘Gastrulation ends’ was taken as the timepoint when contact area between ABplpapppp and ABprppppaa was greater than 0. Intestinal polarized was estimated from visual inspection of the dorsal midline for columnar shaped cells, int5 intercalation^34^ start was decided from lateral views and end from dorsal views, valve bilateral cell shape change was decided from ventral views.

#### Cyst-to-tube transition

Live images were taken every 2 min to catch the dynamics of PKC-3. Unexpectedly, laser power changed only in timepoint 7 for some embryos during multipoint acquisition, and therefore t7 only was multiplied by 1.3 (the ratio of t7/t8 background values) before further analysis. Rolling ball background subtraction (250 pixel radius) was performed for both PKC-3 and *pha-4p*::PM channels and PKC-3 intensity was corrected using single exponential bleach correction^98^. Intensity was summed over time within box placed at the posterior intestine encompassing the entire int8 interface and divided by intensity of initial timepoint. Apical length and interface length in YZ were estimated from PKC-3 and *pha-4p*::PM membrane signal respectively (average of 3 measurements) from resliced, maximum projection images and divided to give percentage coverage of interface. Data for each embryo were aligned to time point when cyst-to-tube transition is complete.

#### Apical ratio in vir ablation embryos

For each embryo with ventral oriented toward the coverslip, PKC-3 intensity at the level of the most ventral end of the int8 nuclei was divided by PKC-3 intensity at the more dorsal intestinal midline (int7/8 junction).

#### HMR-1^nhr-67^(–) quantification

Images were background subtracted using mean of three estimates of the slide background near the embryo inside the eggshell. Two points were manually placed on the lateral membranes of visible vir cell using brightest HMR-1 pixels and/or CAAX signal and intensity was averaged over a 3-pixel-radius ball placed at each point. For plotting, to account for variability in exposure times used on different imaging dates, intensities for each imaging date were divided by the median of the intensities for array negative control embryos. Strong degradation threshold was set at one standard deviation below the mean of array negative HMR-1 intensities.

#### Colocalization of PAR-6^vir^ and PKC-3

A 6-8μm line along end of rectum or intestine in the plane of apical surface where PKC-3 was brightest was used to calculate linescan intensity values measured with imageJ function Plot Profile. A Gaussian fit of PAR-6^vir^ was used to calculate center and the distance in both PAR-6^vir^ and PKC-3 channels was offset. A mean for 12 embryos measured was calculated by first linearly interpolating the linescan values to a common set of distances before plotting the mean and 95% CI of intensities.

#### Puncta intensity quantification

Points were manually placed at the center of the vir-intestine and vir-rectum puncta. If no clear puncta was visible even after adjusting contrast to show dim signals, the brightest pixel near the center of the two gut nuclei was used. Fluorescence signal was averaged over a 3-pixel-radius ball placed at each point for each embryo and the mean of three background measurements outside the embryo was subtracted. To calculate the ‘int-facing puncta’ percentage, the intensity at the intestine-facing puncta was divided by the sum of the intestine-facing and rectal-facing puncta for each embryo. For Figure 3H, ‘rectal-facing’ intensity was shown for clarity and is equivalent to 100 – ‘int-facing’ percentage. To plot the ‘total’, the sum of the intestine-facing and rectal-facing puncta for each embryo was divided by the mean of all control embryos and divided by 2. Individual points for each embryo plotted using swarmplot along with mean and standard deviation of each condition.

### Statistics

Statistics were performed using Excel or scipy.stats^99^. The unit of biological replication was considered one embryo. For comparisons among multiple groups (Figure S1N), one-way ANOVA was used, and if significant, a post-hoc Tukey’s test for pairwise comparisons. For Figure 2F: Fisher’s Exact Test was performed between all control and all HMR-1^nhr-67^(–) animals regardless of degradation strength. For comparisons between two conditions (Figure 2D, Figure 3H, K, Figure 4D, 4F), a either a two-tailed student’s T-test or a nonparametric Mann-WhitneyU test was performed when data did not meet the assumption of normality.

