## Supplementary material for "Reciprocal E-cadherin signaling aligns apical surfaces between neighboring epithelial tissues to complete the *C. elegans* digestive tract": Figures S1-S5

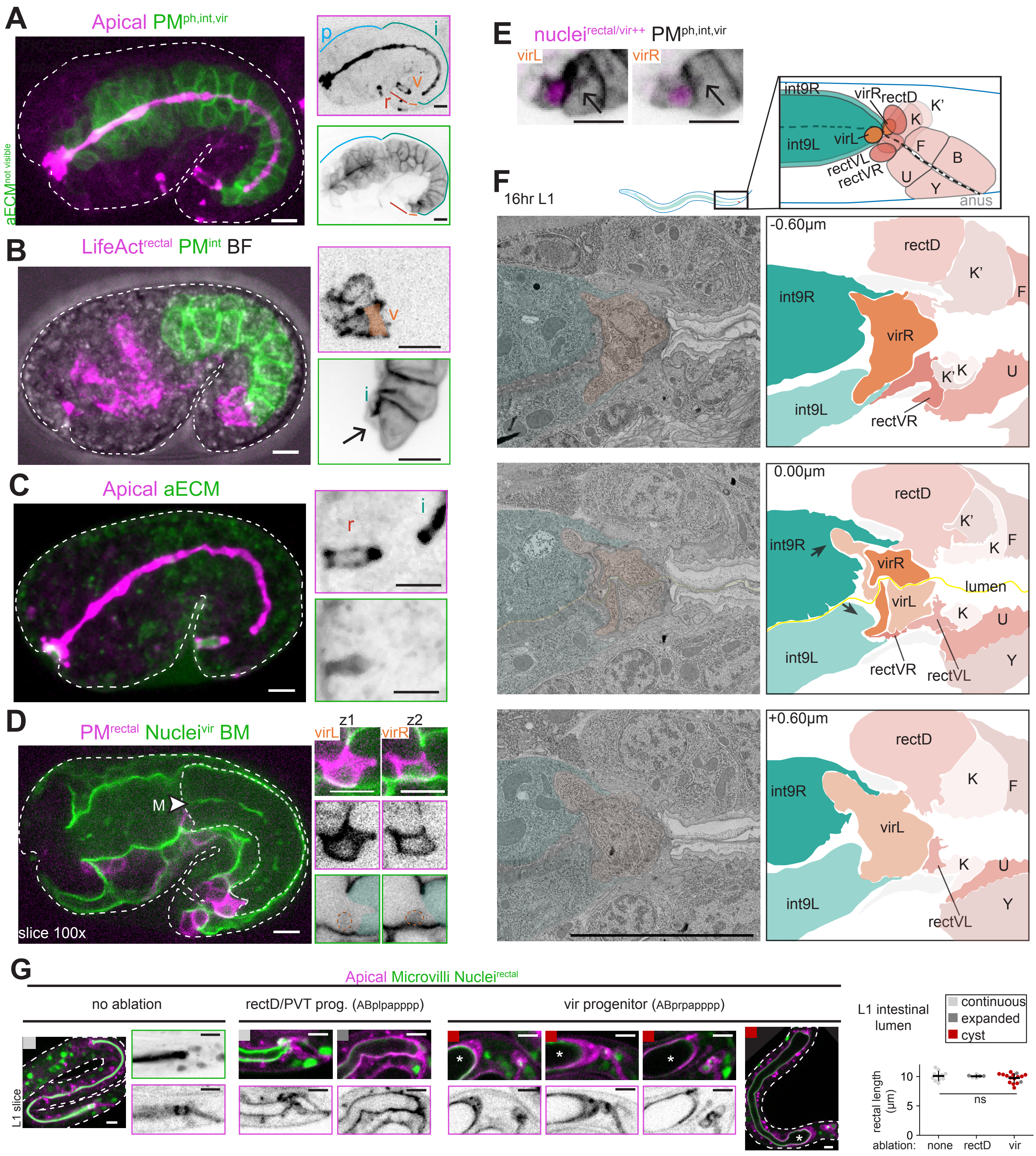


**Figure S1. Rectal valve (vir) cells facilitate the connection between the intestine and rectum**

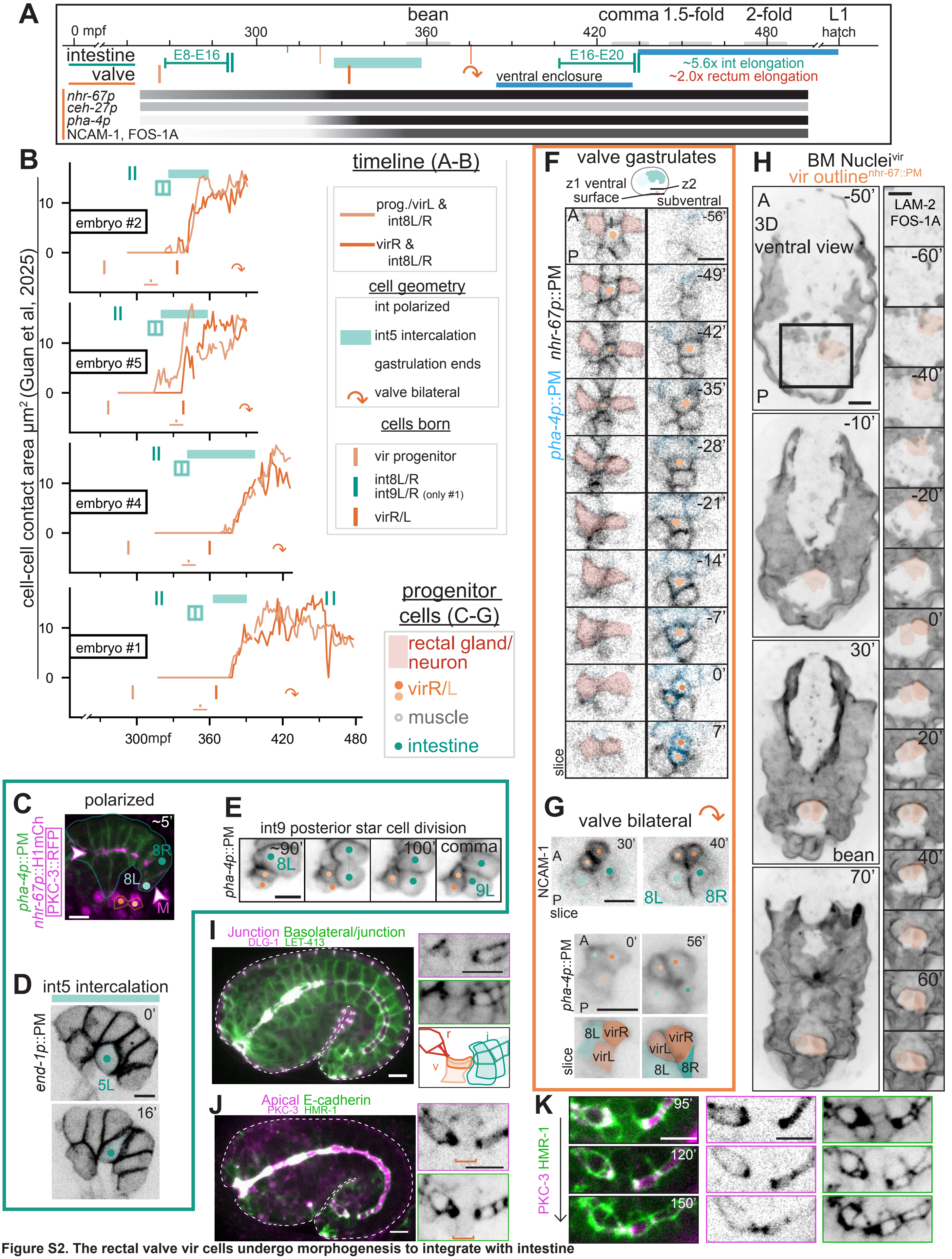


**Figure S2. The rectal valve vir cells undergo morphogenesis to integrate with intestine**

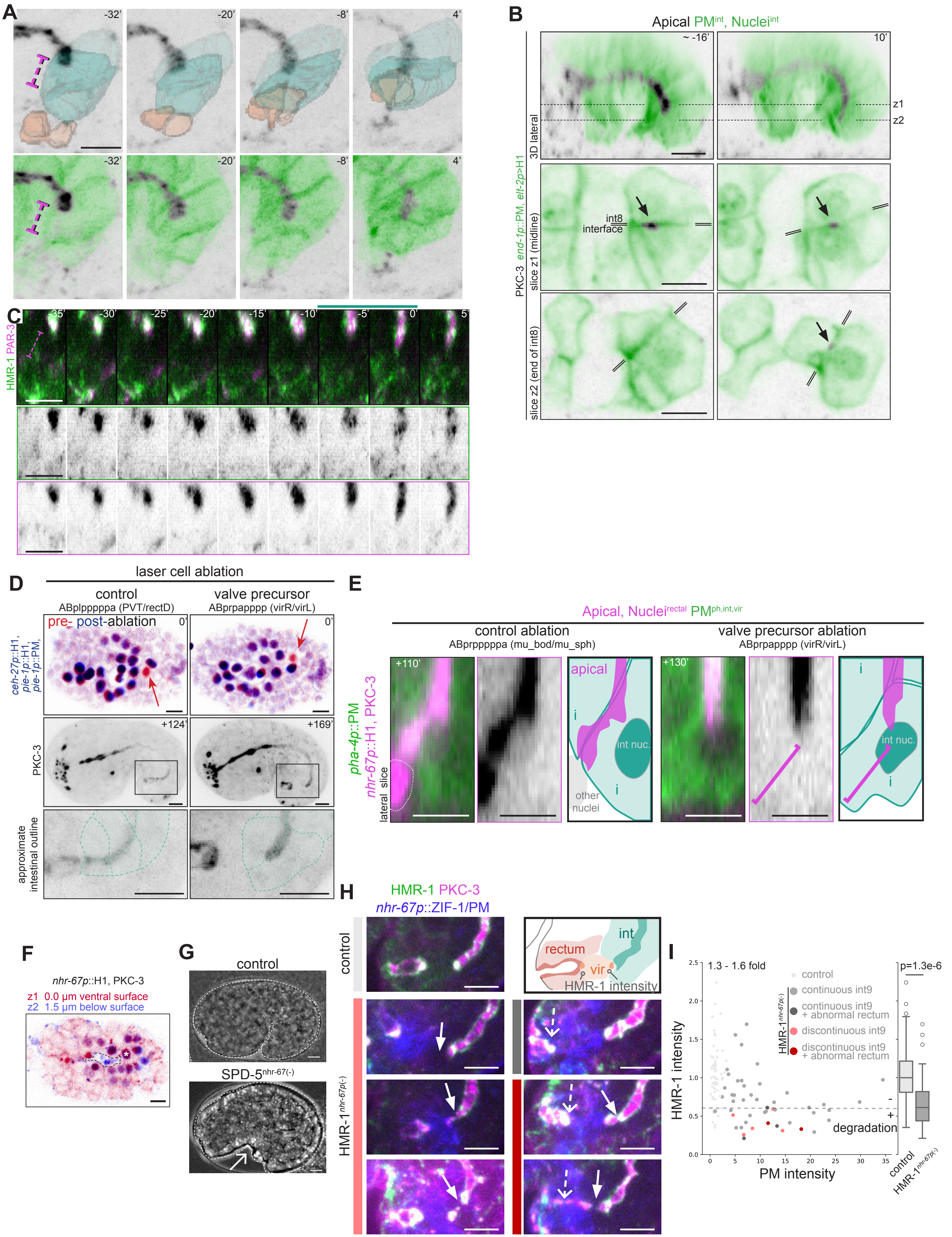


**Figure S3. The intestinal cyst-to-tube transition requires vir cells**

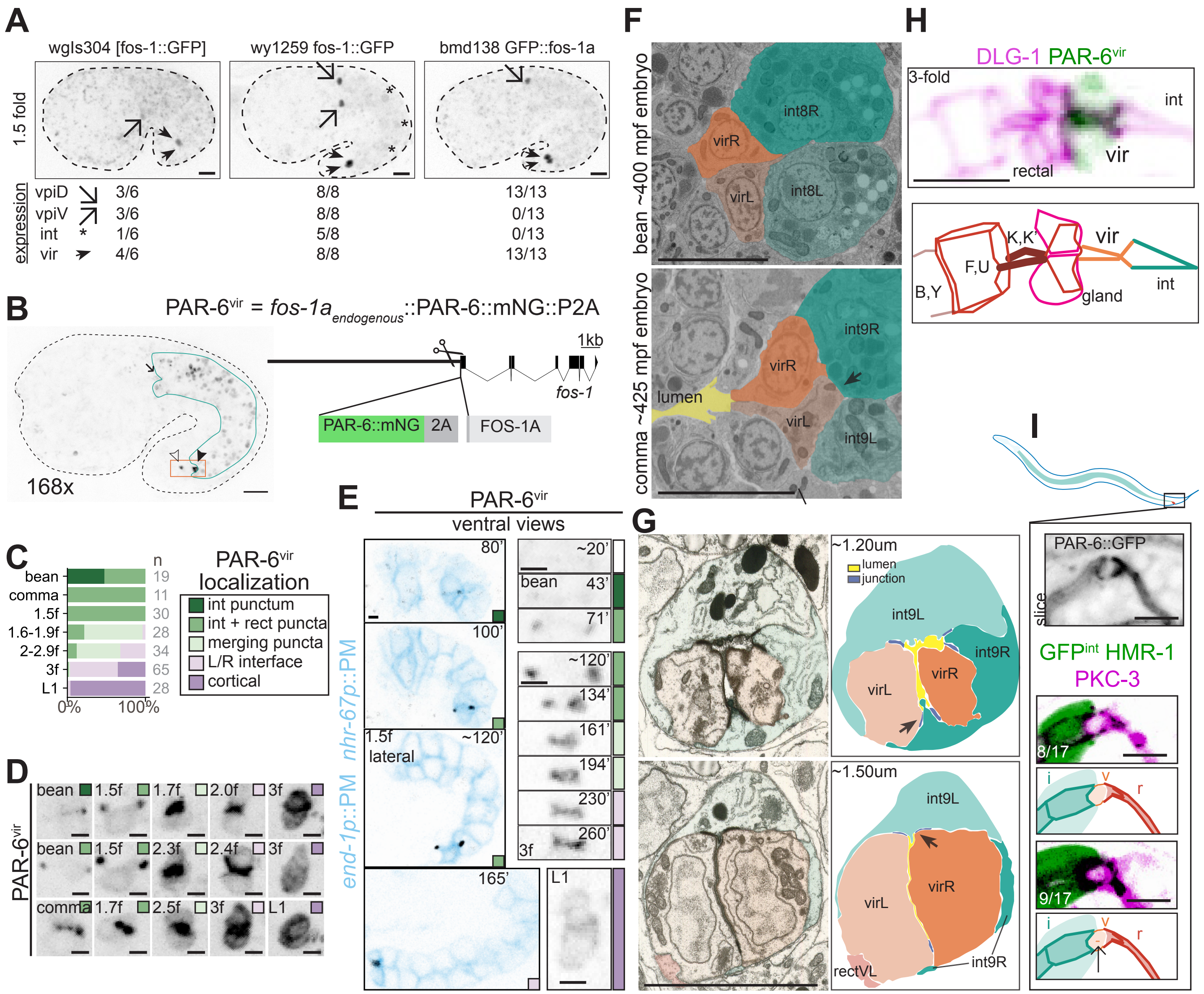


**Figure S4. vir cells are bipolar**

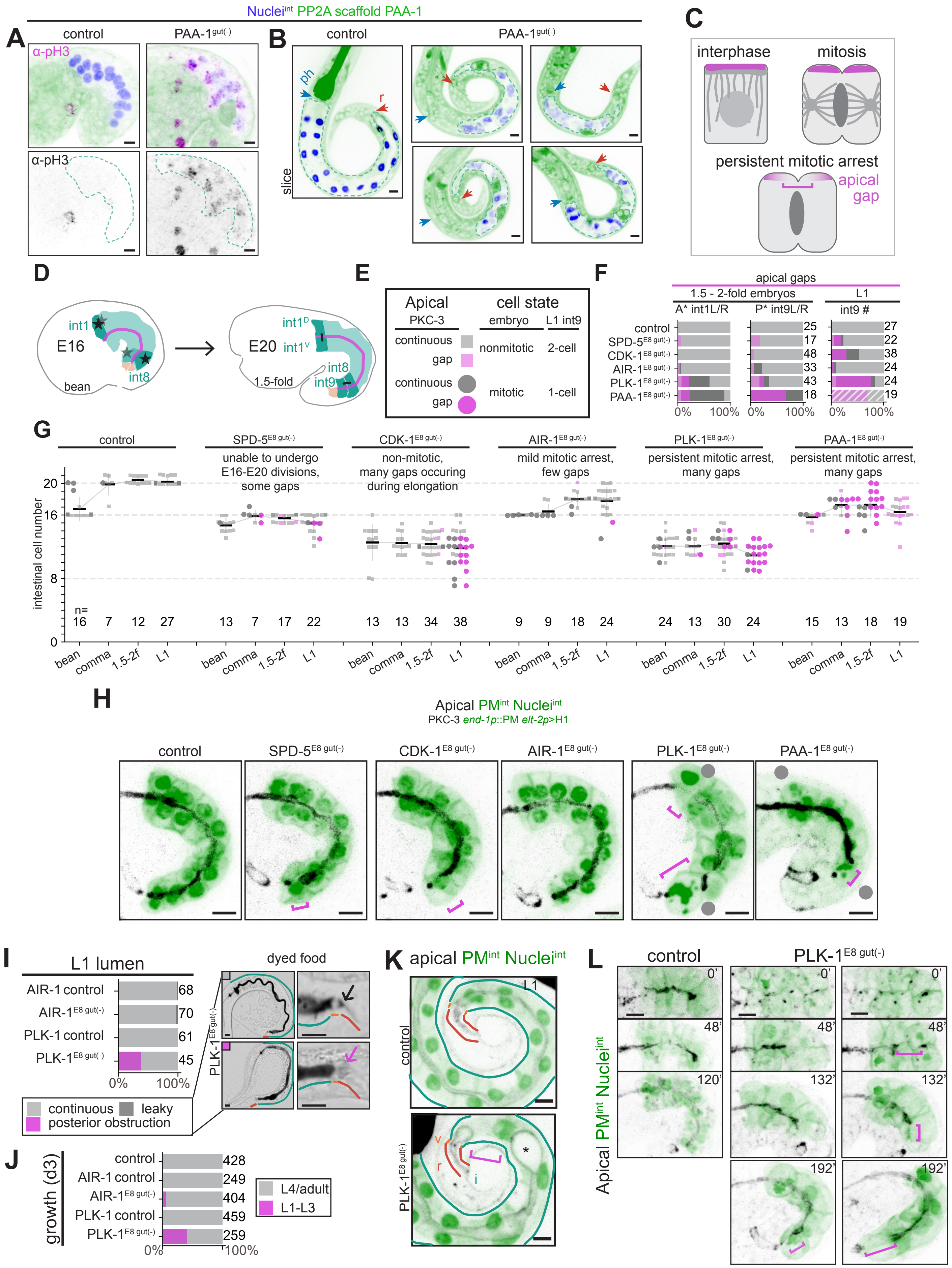


**Figure S5: Consequences of mitotic perdurance in the intestine**
